# Supporting decision making: Modeling and forecasting measles in a London borough

**DOI:** 10.1101/497800

**Authors:** Stefan Edlund, Derryn Lovett, James Kaufman, Kezban Yagci Sokat, Johan van Wijgerden, Alan J. Poots

## Abstract

To investigate the feasibility of using freeware to model and forecast disease on a local scale, we report the results of modeling measles using a spatial patch model centered around 73 clinics in the North West London Borough of Ealing. MMR1 and MM2 immunization data was extracted for three cohorts, age 1-3, 4-6 and 7-19 and patient population was estimated using general practice profile records.

We designed the measles model using the open source Spatiotemporal Epidemiological Modeler (STEM), extending a compartmental disease model to include both maternal immunity and delays in antibody response after immunization. Individuals above age 19 are not included in the modeling.

Next, we generate an approximate 20-year model of vaccination coverage for Ealing. In England, children are immunized between age 1 and 2, then again at around age 5; hence immunization events are modeled for the age 1-3 and age 4-6 cohorts. Parameter values were based on measles research literature; transmission coefficients were estimated using the Polymod contact data and also fitted to 2011-2012 case reporting data for Ealing.

To examine possible effects of policy change, we create two scenarios A and B. In A, we increase vaccination coverage by 10% for all clinics; in B, we focus only on the bottom 10% of the poorest performing clinics (8 clinics total) and equivalently improve their coverage. Scenario A reduces measles from an initial level of 60 cases per year (2011) to 26 cases per year in 2017 (a 58% reduction), compared to the status quo which declines to 45 cases per year. Scenario B reduces measles by 44%, or to 34 cases per year in 2017.

We conclude that local scale modeling is possible, and that the transparency of analysis provided by an open source application lends credence to the output of the models.

## INTRODUCTION

The MMR vaccine is an immunization vaccine against measles, mumps and rubella and consists of live attenuated viruses from all three diseases. In developed countries, the vaccine is usually administered to children around age 1 and again before starting school at about age 5. In high-risk developing countries where healthcare infrastructure is lacking, mass vaccination campaigns have been credited with reducing the global measles disease deaths by 71% between 2000 and 2011 [1].

Yet, even in developed countries, there has been an increase in measles cases; in 2012, there were more than 2,000 cases of measles in England and Wales alone, the highest figure for two decades [2]. The UK epidemic has been blamed in part on misleading information linking diseases on the autism spectrum to the MMR vaccine [3], leading to an overall reduction in the coverage of the vaccine. Estimates of the reproductive number of the virus (*R*_*0*_) show that an increase in *R*_*0*_ occurred almost immediately after the decrease in the MMR vaccine uptake in 1998 [4].

The Spatiotemporal Epidemiological Modeler (STEM) provides an open source framework for building and evaluating spatiotemporal epidemiological models [5]. In addition to textbook examples of disease models, STEM has denominator data for the entire world, including population estimates, administrative regions with common border relationships, air transportation and road transportation networks, and earth science data (e.g. elevation and surface temperature). An open source project under the Eclipse Foundation, STEM can support collaboration within a community; research teams can share models they create as open contributions to STEM that can be freely used and extended thereby supporting peer-to-peer collaboration, review and refinement.

Here we present a localized measles study for the North West London borough of Ealing (Figure 1), to examine the feasibility of using STEM at a high geographical and temporal resolution to support decision-making. STEM is used to model the disease burden of measles and to forecast two scenarios congruent to policy options. As a localized study, both spatially and temporally, we do not expect to reproduce the epidemic that the rest of England and especially Wales have experienced in recent years. Instead, our intent is to assess the potential impact of changes in MMR vaccination coverage for a local set of General Practitioners (GPs) in Ealing; thereby guiding recommendations as to which strategy would have best effect (particularly relevant given limited resources) and testing the feasibility of modeling for local public health decision makers.

**Figure 1.**
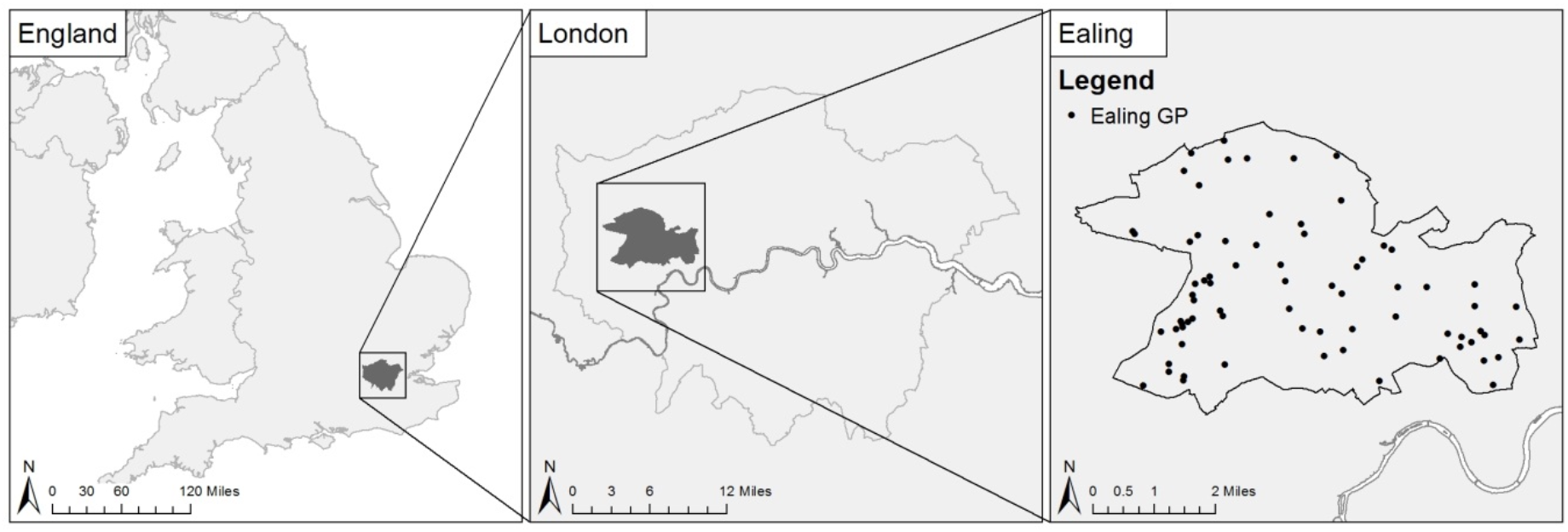
Locations of Ealing and Ealing General Practice Clinics.

## METHODS

### Summary

Time-series data on vaccination coverage between age 1 and 2 (MMR1) and booster vaccination at or around age 5 (MMR2), are used to build a measles model parameterized from measles research literature. We studied four cohorts in particular, age 0-<1, 1-3, 4-6 and 7-19. The second and third cohorts were chosen around the average of the age of vaccination. We cut off the last cohort at age 19; since measles is rare in the adult population, we assume immunity is acquired at this point.

We build two predictive scenarios out 5 years into the future. In the first, scenario A, we improve vaccination coverage for both initial and booster dose by 10% for all clinics, capped at 100%. In the second, scenario B, we pick the 10% worst performing clinics and similarly improve their coverage.

### Demographic and Inoculation Data

Ealing Primary Care Trust (PCT), now with the NHS transformation a Clinical Commissioning Group, provided vaccination data used in our evaluation. The data include two years (2011 and 2012) of monthly MMR1 and MMR2 vaccination coverage for 73 GPs in Ealing. The data were provided as two components: a denominator representing the total number of children having a birthday in a given month for whom the clinic was responsible, and a numerator indicating the number of children having a birthday in the month who received a vaccination. MMR1 coverage data were provided for the age 1-3 cohort; whilst MMR1 and MMR2 coverage data were provided for the age 4-6 and age 7-19 cohorts.

An examination of the vaccination coverage of the 73 Ealing GPs (see supplementary file) is that it has increased from January 2011 to October 2012 (i.e., compare age 1-3 and 4-6 versus 7-19), which could be due to more positive media reporting of MMR vaccination, NHS-led vaccination campaigns and improved data quality as consequence of local improvement initiatives, such as the feedback of GP data to the practices.

### Spatial Model

The spatial model used for the 73 GP clinics was constructed using a Voronoi tessellation method [6] around the geographic coordinates of the clinics (Figure 2). These do not represent the true extents, which are overlapping, but provide a medium for visual representation of change to be displayed. This method is scalable to large numbers of GPs without a burdensome collection and digitization regimen.

**Figure 2.**
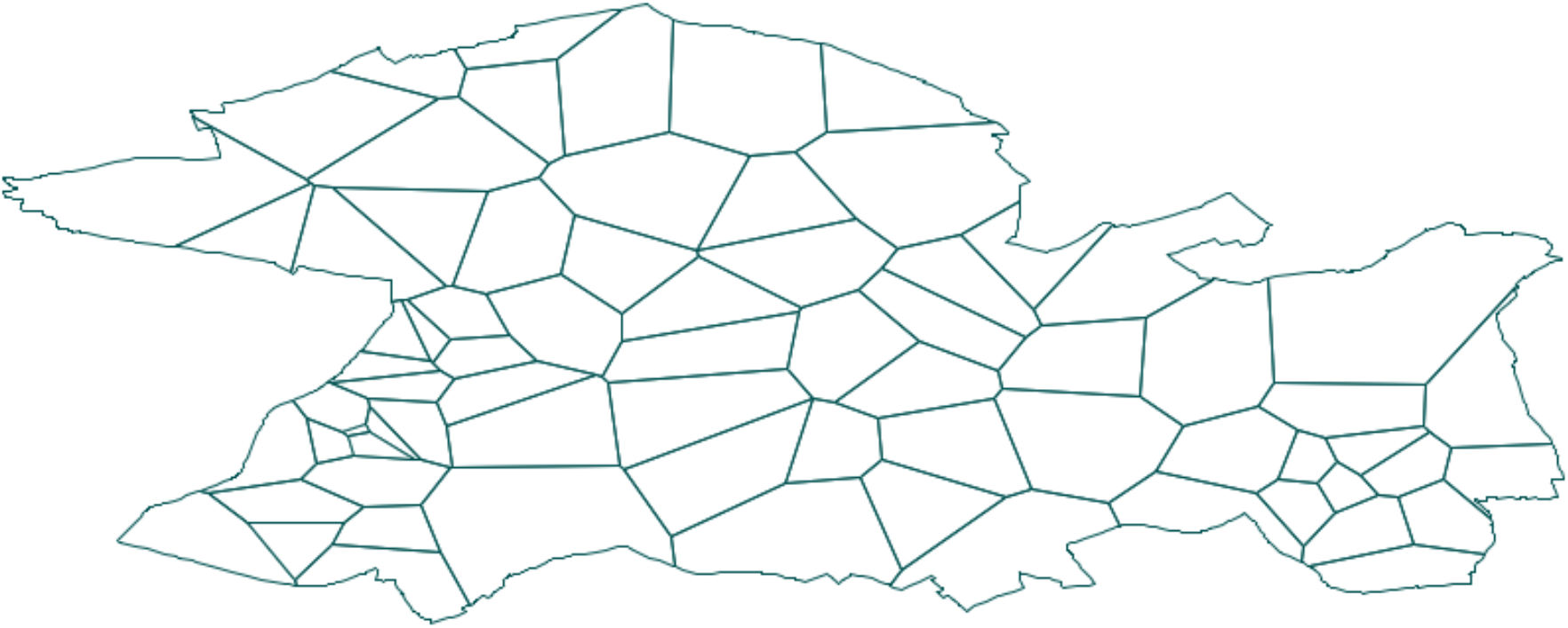
Voronoi Tessellation around the geographic coordinates of the 73 Ealing General Practices.

The total patient population for each clinic was estimated from GP profile records provided by the Association of Public Health Observatories (APHO), and the cohort ages were assumed uniformly distributed. The total population age 0-19 for all of Ealing was estimated at 95,775.

In Figure 3 and Figure 4, we show a plot of vaccination coverage averaged over all 73 clinics in Ealing. For the plot in Figure 3, one data point should be interpreted as the fraction of children having their birthday in the given month and who received one dose of MMR vaccine on or after their first birthday. A data point in Figure 4 represents the fraction of children whose birthday falls within the month and who received two doses of MMR vaccine, where the first dose was given on or after their first birthday.

**Figure 3.**
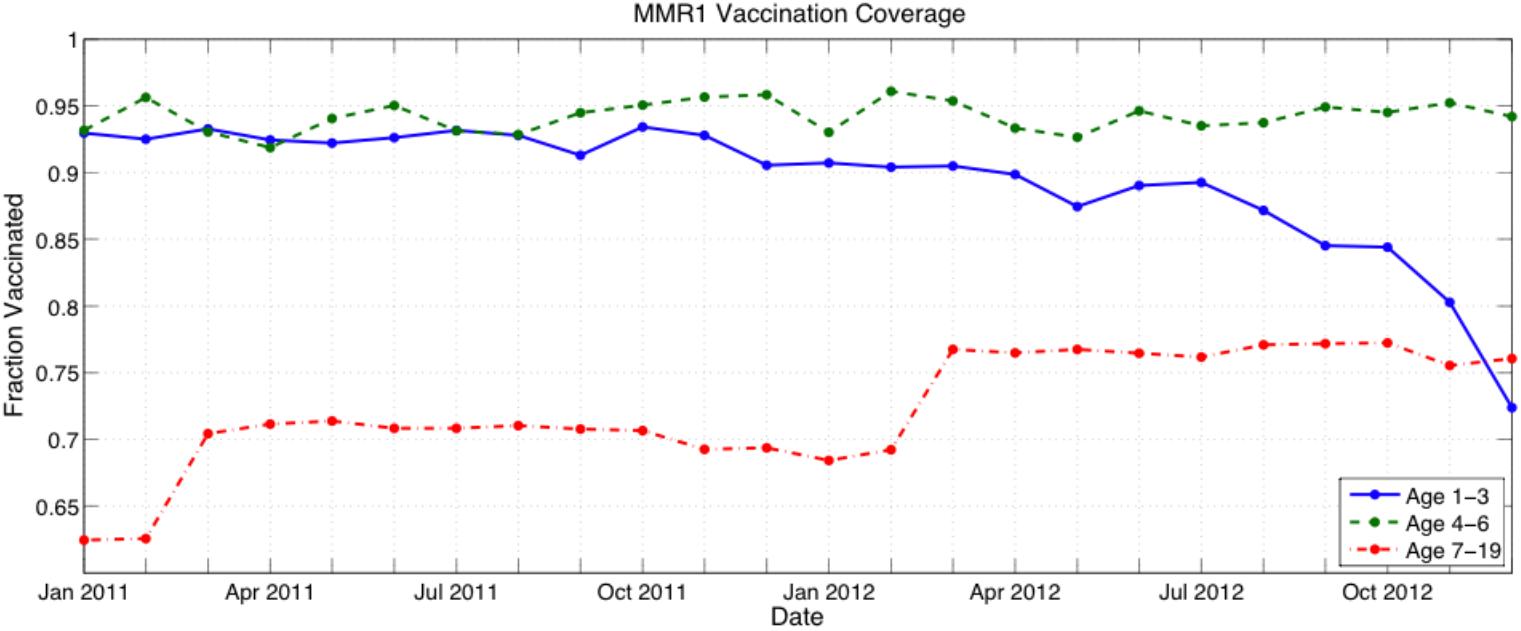
MMR1 Vaccination Coverage for 2011 and 2012. One data point should be interpreted as the fraction of children having their birthday in the given month and who received one dose of MMR vaccine on or after their first birthday.

**Figure 4.**
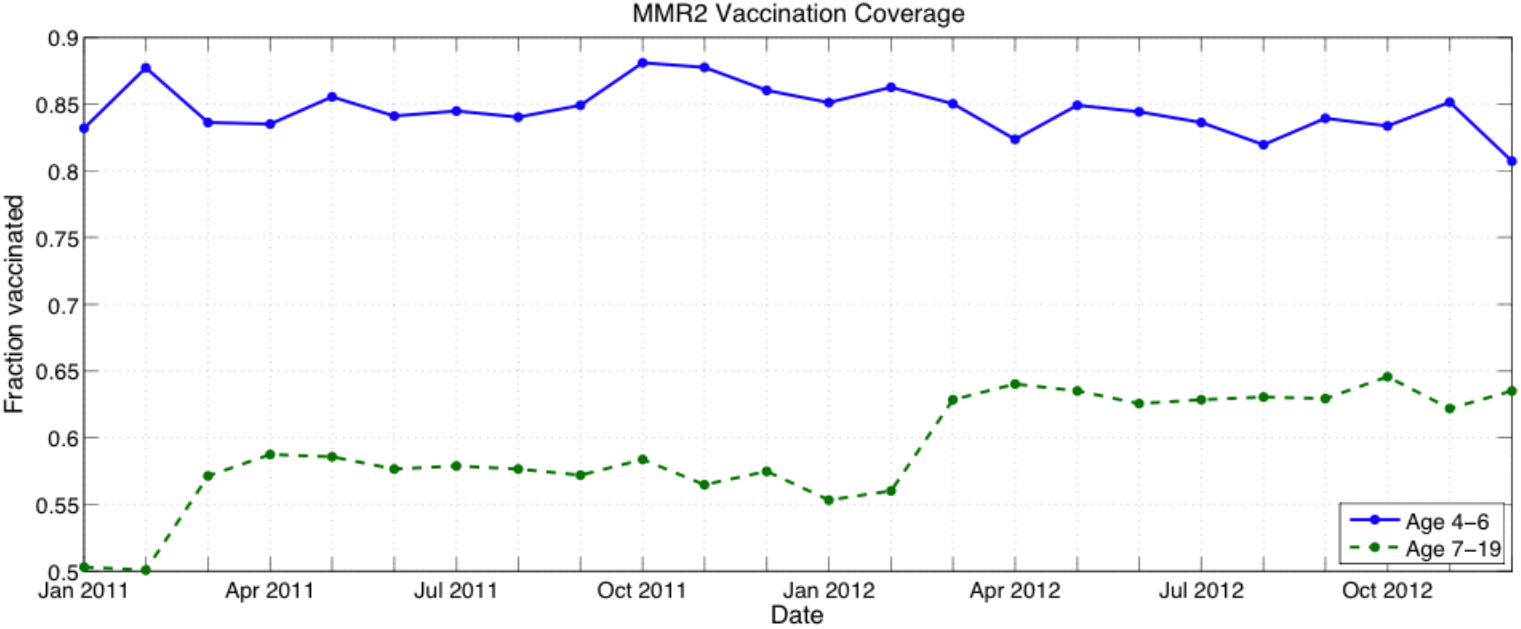
MMR2 Vaccination Coverage for 2011 and 2012. A data point represents the fraction of children whose birthday falls within the month and who received two doses of MMR vaccine, where the first dose was given on or after their first birthday.

**Figure 5.**
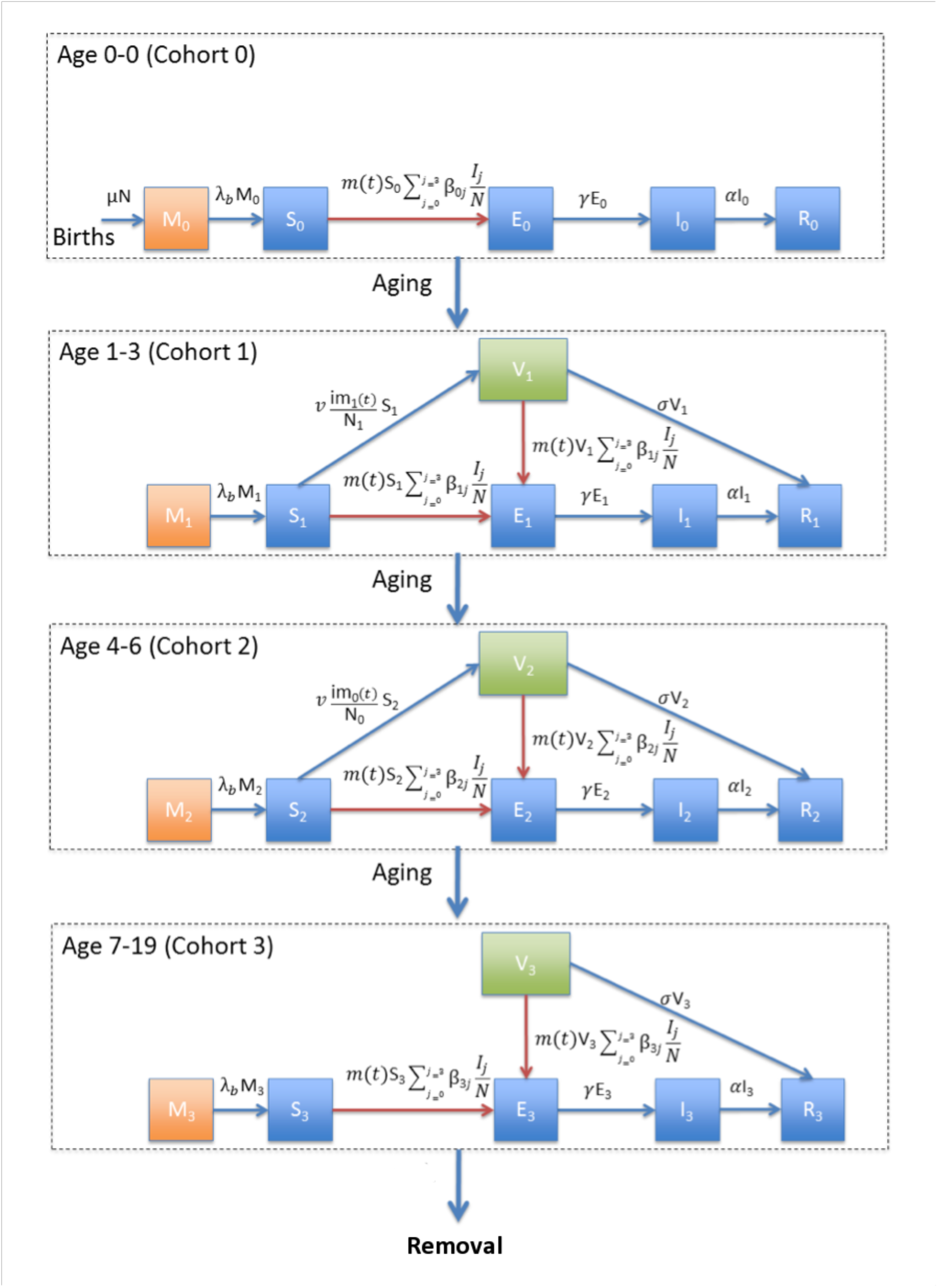
Measles Transmission Model. All four cohorts are modeled separately, and children age out of one cohort into the next over time. After age 19, children are considered immune and removed from the system.

### Measles Transmission Model

A SEIR compartmental model [7] is developed to study the spread of measles and the effect of vaccination, extended with an M compartment to capture maternal immunity and a V compartment for individuals vaccinated (but not yet immune). M, S, E, I, R and V represent the maternal immunity, susceptible, exposed, infected, recovered and vaccinated populations respectively. N is the total population size which is the summation of all children and adolescents in all these states at a given time period (N=M+S+E+I+R+V).

The differential equations of the system are given below. The number in subscript represents the individual cohorts, where 0 is age 0 and 1 is age 1-3 etc. There are four sets of equations for the four cohorts modeled. In the first set of equations (Eq.1a-e), new children are added to the population with a rate of *μ*. Initially, a child is protected by the maternal immunity for a period of (1/*λ*_*b*_) year where *λ*_*b*_ is the maternal immunity loss rate. This is also the rate they enter the susceptible compartment. The next three terms in the Susceptible equation express the number of new infections due to interactions between infected children and adolescents in all three cohorts. Exposed children are removed from this compartment and enter the Infected compartment with an incubation rate of γ (day^-1^). Infected children are removed from this compartment with a recovery rate of α (day^-1^). The Recovered compartment tracks the number of children who acquired natural immunity by recovering from the infection with a rate of α.

Eq. 2a-f and Eq. 3a-f resemble Eq. 1a-e, but also introduce a Vaccination term [8]. The last term in the susceptible rate change equation accounts for vaccination where ν is the vaccine efficacy. The equation for Recovered is extended with a term for children who acquire passive immunity by vaccination with an inoculated immunity rate of σ (day^-1^). Children enter into the Vaccinated compartment with vaccinations during the Susceptible state; they leave this compartment either via Infections, which is expressed with the next three terms in the vaccinated compartment rate change equation, or by acquiring passive immunity. In addition to the number of new infections due to interactions of Susceptible children with the infected children, the number of new infections due to interactions of vaccinated (but not yet immune) children with the infected children is accounted for the rate change in Exposed children. In Eq. 2-4, the first term in each equation represents aging. In addition to aging, adolescents are removed from the system in the last term for equation set 4. So newborns are entered into the system at a rate *μ* in the first cohort and removed from the system at a rate *μ* in the last cohort. This way we ensure that the total population remains constant.

The last equation (Eq. 5) shows the sinusoidal function used to model seasonal forcing, *m(t)* where δ, θ, and ω represents the magnitude of the seasonal variation, phase shift and modulation period. Seasonal changes in measles transmission can been explained by changes in social behavior, for instance, increased contact and thus increased transmission during the school year [9].

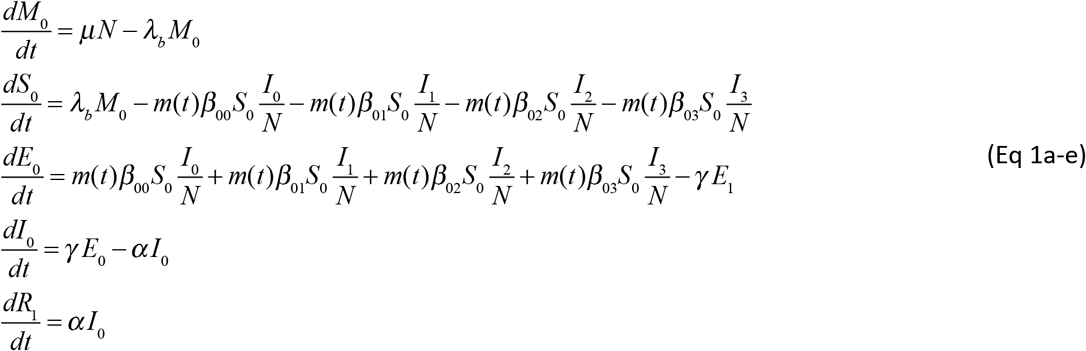

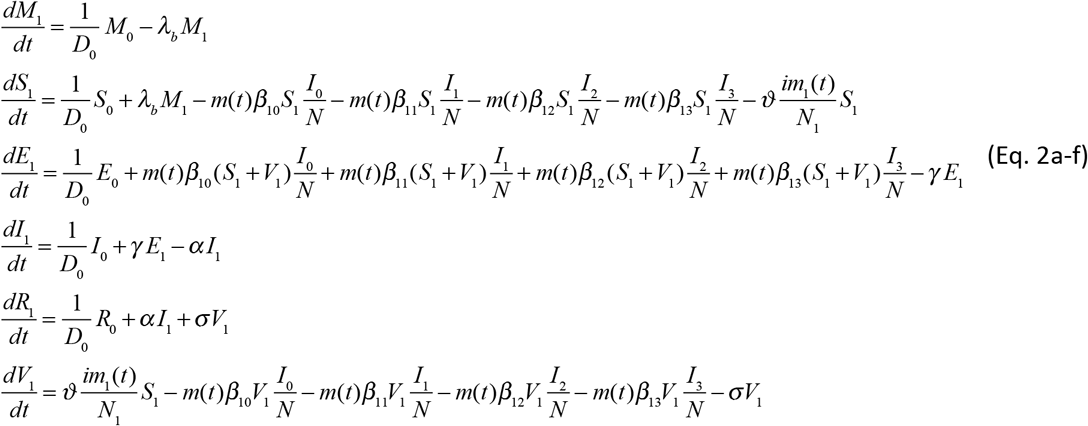

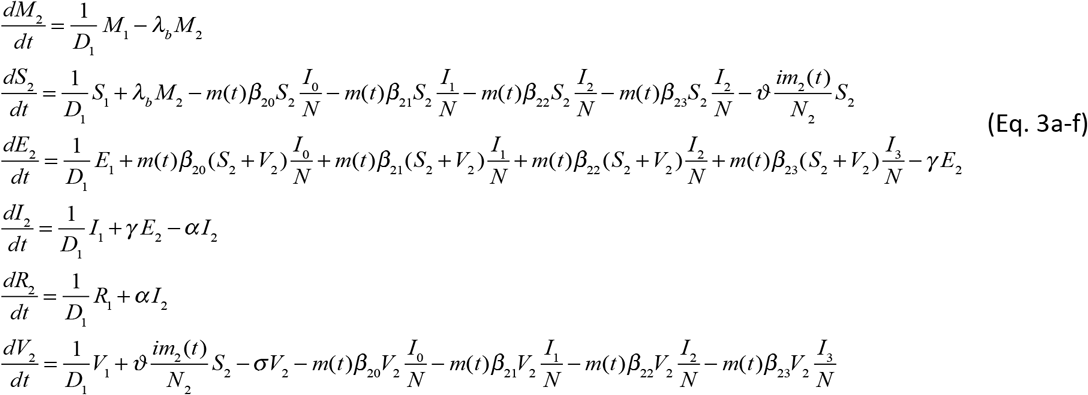

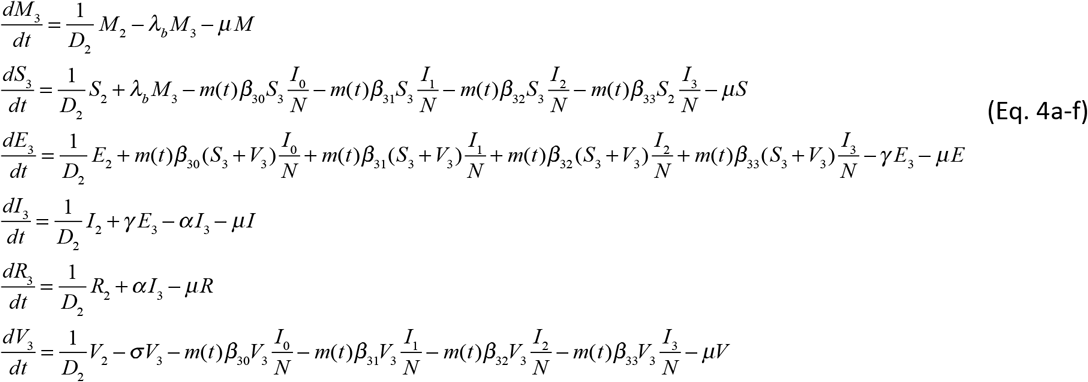

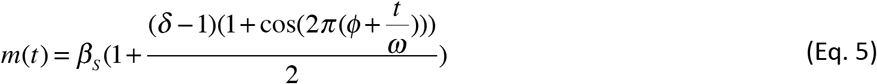

### Parameterization

Descriptions of the parameters, their values and the references are shown in Table 1. The estimated duration of maternal immunity varies from 3 to 6 months (10-13). We use 4 months as advocated by Bjornstad et al. [14].

**Table 1.** Parameter values used in the measles model

| Description | Symbol | Value | Resource/Reference |
| --- | --- | --- | --- |
| Background birth/death rate | $\mu$ | 1/20 (year<sup>-1</sup>) | Constrained by model |
| Maternal immunity loss rate | $\lambda_b$ | 3 (year<sup>-1</sup>) | [a] |
| Transmission rate between cohort i=(0,1,2,3) and cohort j=(0,1,2,3) | $B_{ij}$ | Polymod data | [b] |
| Vaccine efficacy | $v$ | 0.95 | [c] |
| Total MMR 1 immunizations at time t | $im_1(t)$ | Ealing PCT | [d] |
| Total population cohort i =(0,1,2,3) | $N_i$ | APHO/ONS | [e] |
| Incubation rate* | $\gamma$ | 0.1 (day<sup>-1</sup>) (.06-.14) | [f] [g] |
| Recovery rate* | $\alpha$ | 0.125 (day<sup>-1</sup>) (.2-NA) | [h] [i] |
| Inoculated Immunity Rate | $\sigma$ | 0.1 (day<sup>-1</sup>) | [j] |
| Length of cohort 0 | $D_0$ | 1 year | |
| Total MMR 2 immunizations at time t | $im_2(t)$ | Ealing PCT | [d] |
| Length of cohort 1 | $D_1$ | 1 year | Length of cohort 1 |
| Length of cohort 2 | $D_2$ | 3 years | Length of cohort 2 |
| Length of cohort 3 | $D_3$ | 3 years | Length of cohort 3 |
| Length of cohort 4 | $D_4$ | 13 years | Length of cohort 4 |
| Total populations all cohorts | N, M, S, E, I, R, V | APHO/ONS | [e] |
| Magnitude of seasonal variation | $\delta$ | 1.25 | [k] |
| Phase shift | $\theta$ | 0 | |
| Modulation period | $\omega$ | 1 year | |
| Transmission Rate Scaling | $\beta_s$ | 0.432 | Fitted |

\* The values in parentheses represent the minimum and maximum value for the parameter.

The incubation period varies from 7 to 18 days [15]. The most common value of the incubation period is 10 days hence our incubation rate is 0.1 (day^-1^) using the Public Health Agency of Canada [16]. Similarly, the most commonly reported value of infectious period (8 days) is used to obtain the recovery rate [17].

To find the contact rates between cohorts, we use contact data from the Polymod project [18], see Figure 6. The data was compiled from 7,290 participants reporting over 90,000 contacts with different individuals during a day, including age, sex, location and duration. The study indicates that 5- to 19-year-olds are expected to suffer the highest incidence during the initial epidemic phase infection transmitted through social contacts, which can also be seen in Figure 6 where the data for UK is shown.

**Figure 6.**
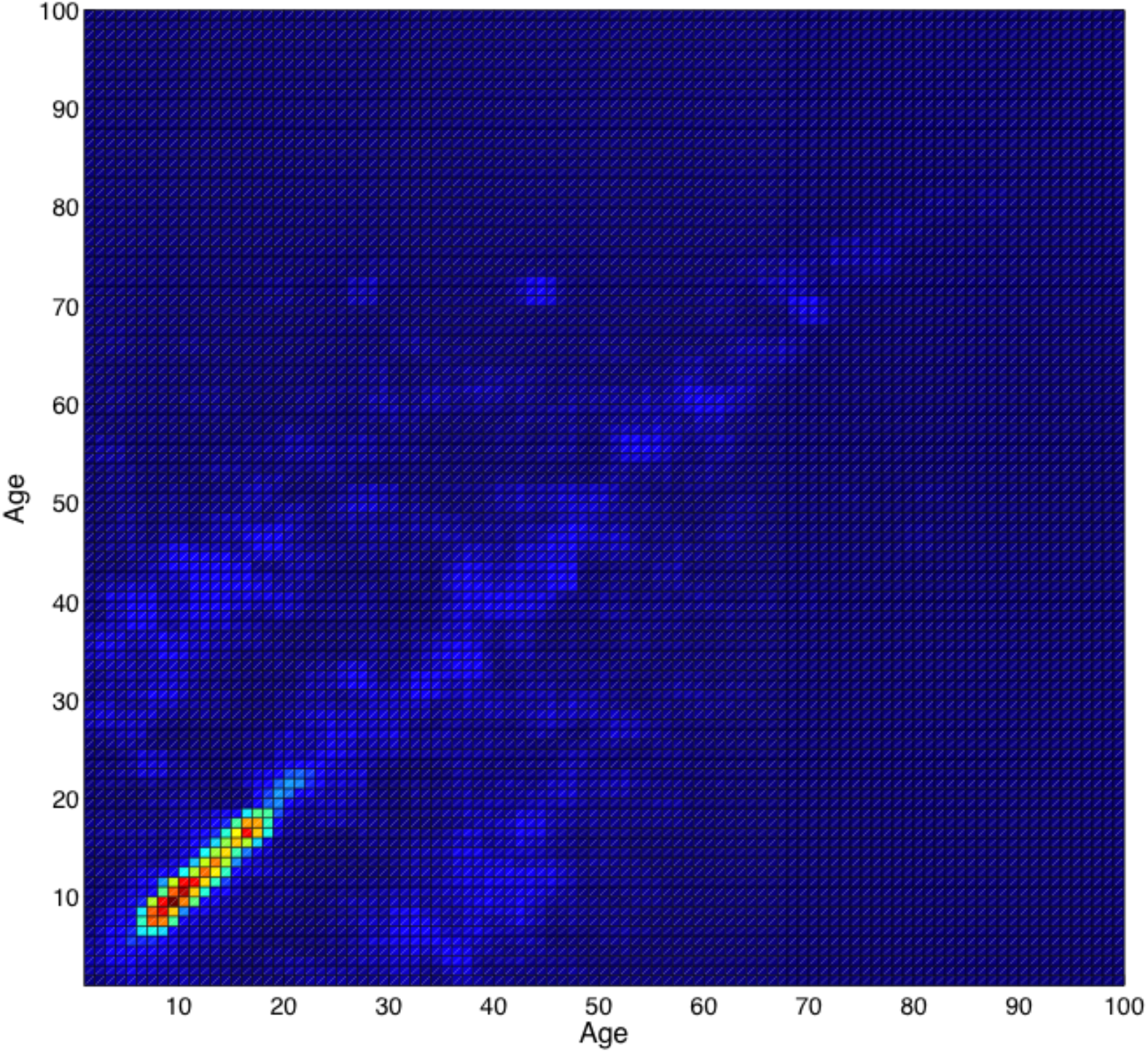
Smooth Contact Matrix for UK. Based on data from Polymod survey as reported by Mossong J et al. (2008), Social contacts and mixing patterns relevant to the spread of infectious diseases, PLoS Med 5(3):e74. doi:10.1371/journal.pmed.0050074

When a child receives a dose of measles vaccine, it takes on average 10 days for the vaccine to give protection [19]. Thus, the inoculated immunity rate σ is 0.1 (day^-1^). Measles vaccine is assumed 95% effective after a dose is applied as per Centers for Disease Control and Prevention [20]. Measles transmission is assumed seasonal, captured as a sinusoidal function using a 1 year modulation period with 25% increase in transmission during peak season [21]. An individual is expected to shed the measles virus for an average of 10 days [17,22].

### Analytical Approach

The first step in the analytic study is to estimate, for each cohort, an initial state of measles in the population. To accomplish this we used the oldest available vaccination coverage data for children aged 7-19 in 2011 and 2012, and ran a model with a random initial condition repeatedly until convergence. We determined that the state of the population at convergence had only a small sensitivity to the initial condition. In particular, over 100 separate runs with randomly set initial condition and after 100,000 days of simulation time, the total incidence (all cohorts) in the last year was 558.2 with a standard deviation of 13.9, within the error tolerance of the integrator. In essence, the oldest vaccination coverage data provided is assumed to have been in place since “beginning of time”. However, the error introduced by this assumption is somewhat mitigated by the fact that adults above age 20 (whose immune state is more difficult to estimate when lacking historic data) are not included in the modeling. STEM was used to run the simulations required to establish the initial conditions.

### Predictive Model

Two different vaccination improvement scenarios were studied for consideration by Ealing public health. For the first strategy (scenario A), we assume improved vaccination coverage by 10% in all clinics (so a clinic with say 80% coverage increases to 88%). In the second strategy (scenario B), only the 10% of poorest performing clinics (8 clinics total) were assumed to increase their vaccination coverage by 10%. These 8 poorest performing clinics were selected according to their vaccination coverage averages for both MMR1 and MMR2 and across all cohorts (except for cohort 0). Measles incidence is modeled 5 years into the future for both of these alternatives, and compared to the status quo where the vaccination coverage is kept identical to the most recent data (MMR1 from age group 1-3 and MMR2 from age group 4-6).

## RESULTS

In Figure 7 panel A, we show the infectious cases on Feb 1 2012 (deeper red indicates more measles cases). This is the reference used in panel B-D. In panel B, the ratio between the number of infectious cases on Feb 1 2017 and the reference is computed. The color legend in the center indicates the computed ratio. In panels C and D, the same ratio is computed for scenario A and scenario B respectively.

**Figure 7A-D.**
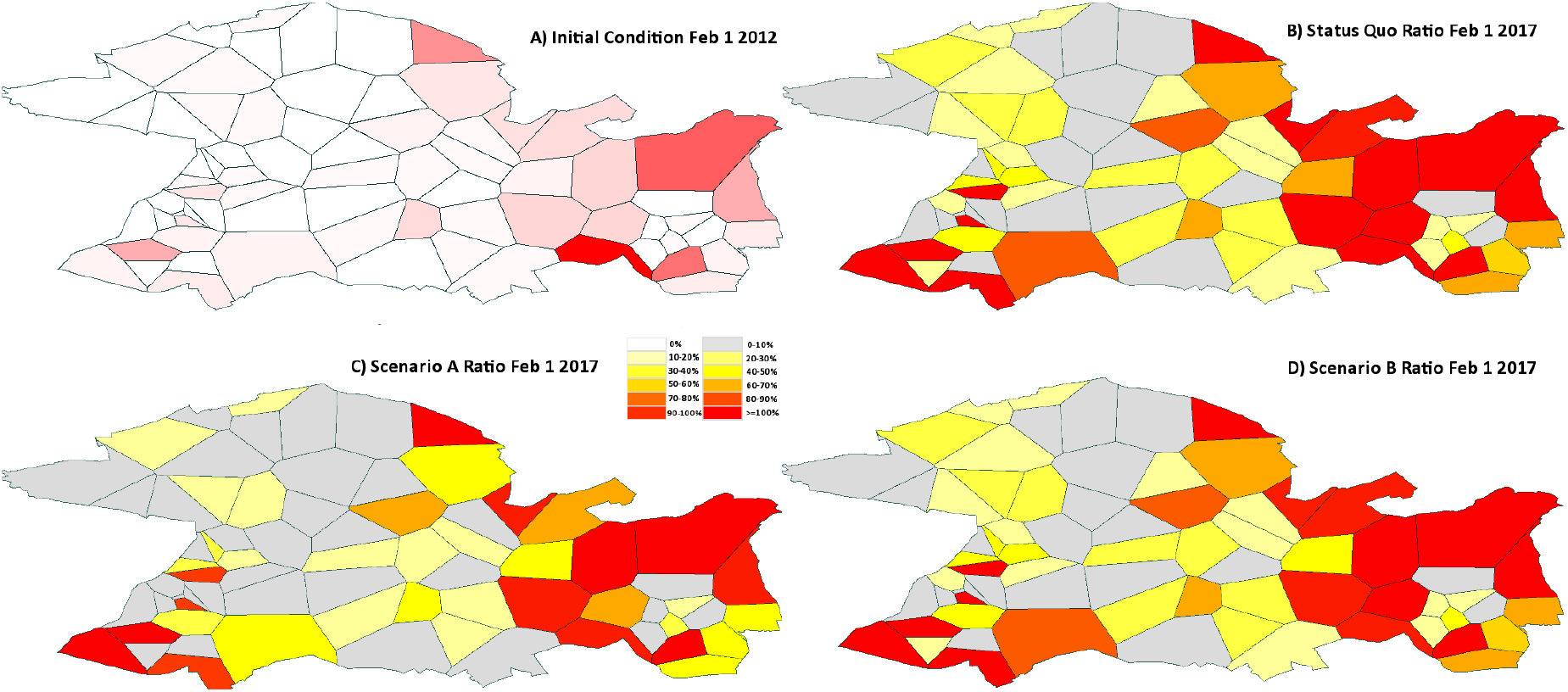
Panel A shows the infectious cases on Feb 2012 (deeper red means more measles cases). Panels B-D show the ratio between the new number of infectious cases on Feb 1 2017 and the reference for status quo, scenario A and scenario B respectively.

In Figure 8 we show the daily measles incidence for the four cohorts at year 2011 and 2012. Total incidence for age 0-19 in 2011 is around 60 cases, dropping to about 42 cases in 2012. In data provided by Public Health England (PHE) measles surveillance program, the total number of cases of measles in 2011 in Ealing was 42 (confirmed + possible + probable); in 2012 the number was 28. This gives us a reporting fraction of about 66-70%, which is within the expected range [23].

**Figure 8.**
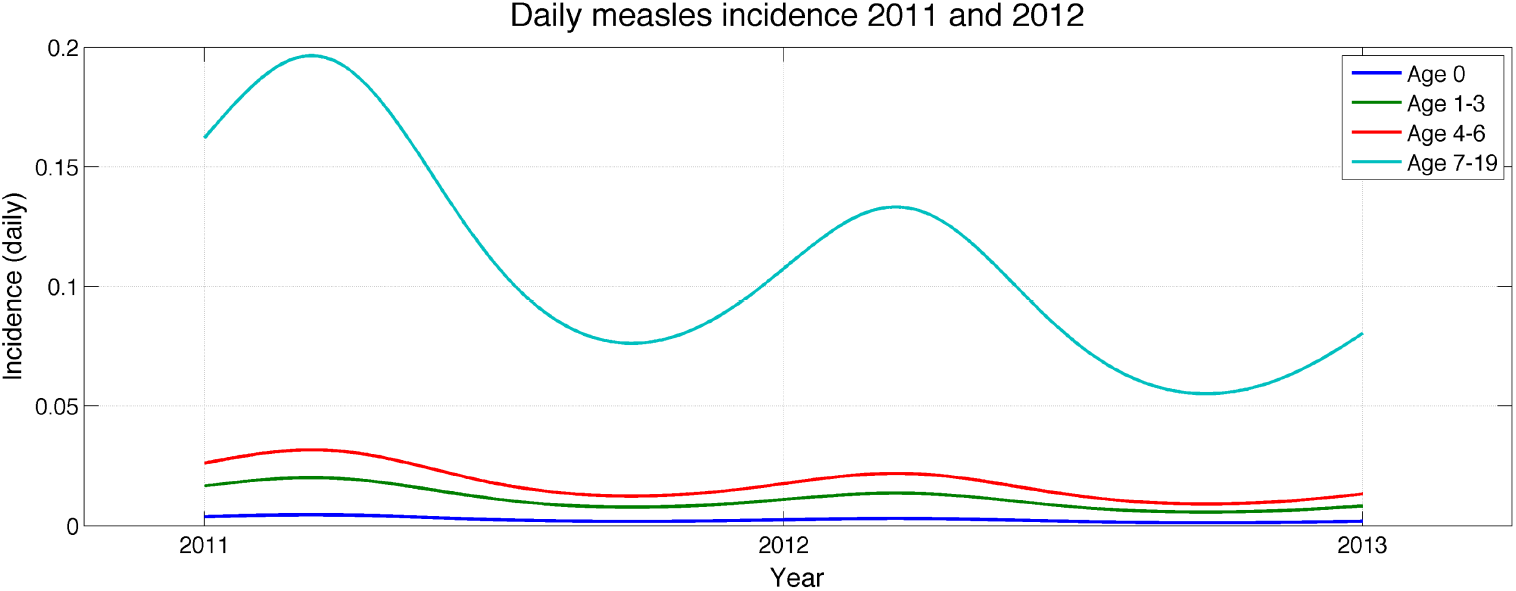
Incidence reported out of the measles model for year 2011 and 2012.

Figure 9 shows total measles incidence predicted out 5 years (2013-2017) for the two scenarios studied as well as the status quo case. When we improve vaccination coverage by 10% for all clinics, measles declines from an initial level of 60 cases per year (2011) to 26 cases per year in 2017, compared to a business as usual scenario in which measles decline to 45 cases per year. When we only focus on improving performance for the bottom 10% of poorly performing clinics (8 clinics total), measles incidence is reduced to 34 cases per year in 2017.

**Figure 9.**
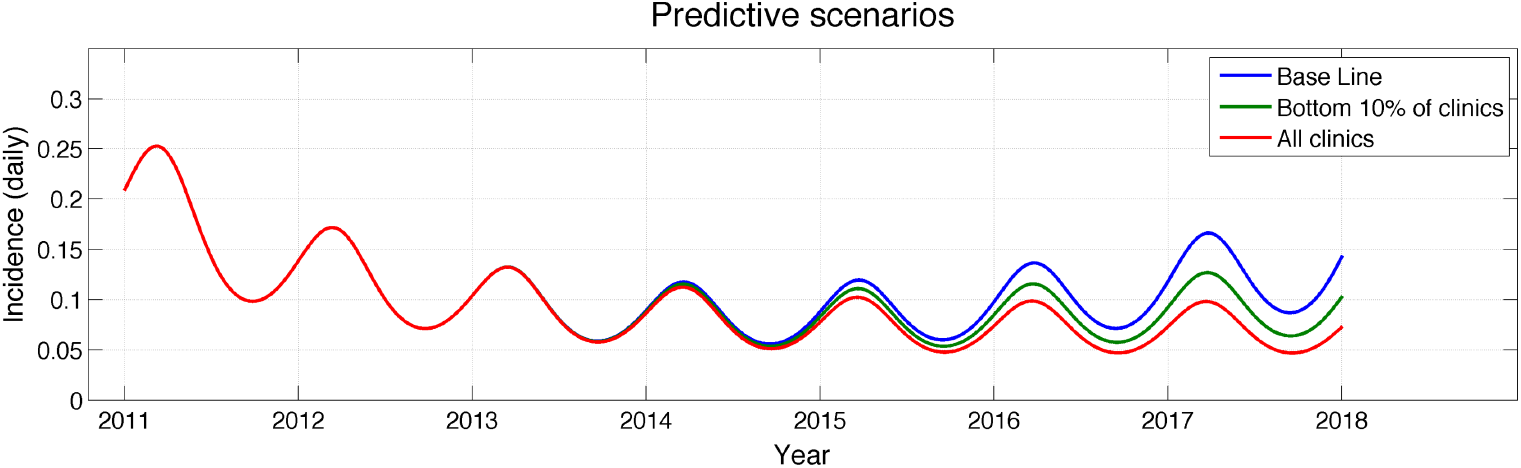
Predictive Scenarios. The plot shows total measles incidence predicted out 5 years (2013-2017) for the two scenarios studied as well as the status quo case.

## DISCUSSION

Our work demonstrates how a tool such as STEM can be used by epidemiologists and public health experts to evaluate the impact of strategies to improve vaccination efficacy. Once a reference model is developed based on available data, it is fairly straightforward to integrate the reference model into the future subject to a range of plausible assumptions. With no intervention or campaign to improve vaccination coverage, the base model predicts a 25% decline in measles incidence. If vaccination coverage is improved by 10% for only the 10% poorest performing clinics (8 of the 73), the model predicts that a relatively large reduction in measles (44%) can be realized over 5 years. Improving the performance of all clinics would yield a 58% decrease in incidence after 5 years.

Selection of an appropriate strategy depends, of course, on economic considerations as well as questions concerning detailed requirements of an education campaign. For example, could the same education materials designed to target the neighborhoods with the lowest vaccination coverage be used for all clinics, or are there specific education needs based on demographic differences between local communities that would require more customized vaccination education campaigns? These questions are beyond the scope of the existing models, but the two alternative intervention models developed here demonstrate how a range of alternative interventions strategies can be quickly evaluated.

### Limitations

There are several challenges to tackle modeling the effect of immunization on measles transmission for a spatially local region such as Ealing.

First, the MMR1 and MMR2 immunization data itself is not perfect. The denominator component of the data (number of children a clinic is responsible for) tends to be too high since tracking children in a borough with a high throughput is challenging. There is often a delay before a child is removed from the register of a clinic when moving out. The numerator (number of children vaccinated) tends to be too low as the vaccination Read-code (the code which the MMR vaccination is recorded against the child) is not always adhered to. GP clinics do not always record full immunization histories on their IT system, particularly if the child is over the age of 5. Consequently, our vaccination coverage data tends to be underestimated resulting in possible overestimation of disease transmission.

Second, the specified measles model itself is not a perfect representation of the world, as with all models. One important factor missing is modeling of imported measles cases from regions external to Ealing. It is possible that some cases are imported due to infected individuals visiting Ealing.

Third, as discussed above, there are challenges in defining the initial condition for our simulation. There is a weakness in the assumption that the oldest MMR1 and MMR2 data we have accurately reflect the performance of clinics going back to the “beginning of time”. Measles vaccination was introduced in the UK only in 1968, but there is no continuous high resolution comprehensive data set back to that time. However, by not considering people born before 1992 in our modeling, we somewhat mitigate the error in that the immune state of older (removed) individuals is ignored. There is also a weakness introduced by the assumption of constant birth rate and death rate, resulting in a uniform distribution of population members into the cohorts used. By taking advantage of historical births and deaths data for Ealing (or if missing, at least UK data), a better estimate of the total size in each cohort could be established. With the promise of electronic health records, over time recording a more comprehensive history of measles in the population should greatly improve our knowledge of the population state and aid in ongoing modeling efforts.

### Strengths

We demonstrated the usefulness of using an open source tool, in this case STEM, both to model infectious disease spread and to measure the impact of alternative intervention strategies such as improved vaccination coverage. The model used is available to any researcher to use freely, allowing transparency of analysis for peer refinement and critique. Furthermore, the new model generator tool available in STEM 2.0 enables even non-expert users to create, build upon and test any model of disease, including the measles model.

Using the shapefile import feature in STEM, custom spatial models can be imported and used directly in STEM, in this case Voronoi polygons derived from locations of medical clinics, allowing easy integration with external tools such as ESRI’s ArcGIS. It is also easy to import time series of vaccination coverage data into STEM, and using a simple drag-and-drop interface to drag those coverage data into any model of disease being studied. This flexibility is important, as measles reporting (and public health reporting in general) is not always based on administrative boundaries or postal codes. Using Voronoi tessellation it was possible to generate a spatial graph based on clinical regions or point of care.

George E. P. Box [24] observed that “essentially, all models are wrong, but some are useful”. Modeling can advise the development of public health policy, but given the uncertainties associated with public health data, it is essential that the assumptions built into such models and the models themselves be fully transparent. Perhaps the greatest strength of STEM is not the use of advanced software technology but the transparency that comes with open source. The Eclipse Foundation provides a community with the tools required so others can build upon existing models to explore the *range of likely outcomes* expected from available data as opposed to running closed or proprietary models generating predictions from a “black box”.

## CONCLUSION

We have demonstrated the feasibility of using open source software (STEM) at fine geospatial and temporal resolutions, with routine data, to provide a resource to support for public health decision makers in examining possible effects of policy change. In the future, we would like to extend the work to a larger spatial region, perhaps all of London or even England and also include imported measles into the model. We would like to investigate the role movement of individuals plays in the spread of measles. In addition, evaluation of a broader range of interventions would be useful; targeting vaccinations to different age groups. Also, we’re interested in modeling the effects on the wider healthcare system, e.g., reductions in emergency admissions and attribution of costs of intervention and disease. Ultimately, the goal is to allow non-technical users to build models of infectious disease, upload data and evaluate interventions strategies for patient benefit.

## Acknowledgements

Thanks to NIHR CLAHRC NWL, NHS and Ealing PCT for sharing the immunization data used in this paper and PHE for providing case reporting data. Thanks also go to Eclipse for its support of the STEM Project.

## Disclaimer

This article presents independent research commissioned by the National Institute for Health Research (NIHR) under the Collaborations for Leadership in Applied Health Research and Care (CLAHRC) programme for North West London. The views expressed in this publication are those of the author(s) and not necessarily those of the NHS, the NIHR or the Department of Health.

## References

1. World Health Organization (2013) Measles fact sheet 286. Available: http://www.who.int/mediacentre/factsheets/fs286/en/. Accessed: 15 December 2013.

2. British Broadcasting Corporation (2013) Measles outbreak in maps and graphics. Available: http://www.bbc.co.uk/news/health-22277186. Accessed: 15 December 2013.

3. Wakefield AJ, Murch SH, Anthony A, Linnel J, Casston DM, et al. (1998) Lleal-lymphoid-nodular hyperplasia, non-specific colitis, and pervasive development disorder in children. Lancet Journal 351(9103): 637-641. RETRACTED.

4. Jansen VAA, Stollenwerk N, Jensen HJ, Ramsay ME, Edmunds WJ et al. (2003) Measles outbreaks in a population with declining vaccine uptake. Science 301(5634): 804.

5. Kaufman J, Edlund S, Douglas J. (2009). “Infectious disease modeling: creating a community to respond to biological threats.” Statistical Communications in Infectious Diseases, Vol 1, Issue 1, Article 1. The Berkeley Electronic Press.

6. Voronoi G (1908). Nouvelles applications des paramètres continus à la théorie des formes quadratiques. Jdie Reine und Angewandte Mathematik 133: 97–178.

7. Anderson RM, May R (1991) Infectious diseases of humans: dynamics and control. New York: Oxford Science Publications.

8. Health Protection Agency of UK (2010) HPA National Measles Guidelines: Local & Regional Services. Available: http://www.hpa.org.uk/webc/HPAwebFile/HPAweb_C/1274088429847. Accessed: 14 December 2013.

9. Wallinga J, Heijne JCM, Kretzschmar M (2005) A measles epidemic threshold in a highly vaccinated population. PLoS Med 2(11): e316

10. Babad HR, Nokes DJ, Gay NJ, Miller E, Morgan-Capner P, Anderson RM (1995) Predicting the impact of measles vaccination in England and Wales: model validation and analysis of policy options. Epidemiol Infect 114:319–41.

11. Gay NJ, Hesketh LM, Morgan-Capner P, Miller E (1995) Interpretation of serological surveillance data for measles using mathematical models: implications for vaccine strategy. Epidemiol Infect 115:139–56.

12. Edmunds WJ, Gay NJ, Kretzschmar M, Pebody RG, Wachmann H (2000) European Seroepidemiology Network. The pre-vaccination epidemiology of measles, mumps and rubella in Europe: implications for modelling studies. Epidemiol Infect 125: 635-50.

13. Simons E, Mort M, Dabbagh A, Strebel P, Wolfson L (2011) Strategic planning for measles control: using data to inform optimal vaccination strategies. J Infect Dis 204(Suppl 1):S28–34.

14. Bjornstad ON, Finkenstadt BF, Grenfell BT (2002) Dynamics of measles epidemics: estimating scaling of transmission rates using a time series SIR model. Ecol Monogr 72(2): 185-202.

15. Health Protection Agency of UK (2010) HPA National Measles Guidelines: Local & Regional Services. Available: http://www.hpa.org.uk/webc/HPAwebFile/HPAweb_C/1274088429847. Accessed: 14 December 2013.

16. Public Health Agency of Canada (2013) Measles. Available: http://www.phacaspc.gc.ca/im/vpd-mev/measles-rougeole-eng.php. Accessed: 14 December 2013.

17. Perez L, Dragicevic S (2009) An agent-based approach for modeling dynamics of contagious disease spread. Int J Health Geogr 8:50.

18. Mossong J, Hens N, Jit M, Beutels P, Auranen K, et al. (2008) Social contacts and mixing patterns relevant to the spread of infectious diseases. PLoS Med 5(3): e74. doi:10.1371/journal.pmed.0050074

19. The MMR vaccinae and your questions answered. Available: http://www.babyexpert.com/baby/health/the-mmr-vaccine-and150-your-questions-answered/2207.html. Accessed: 9 January 2014

20. Fiebelkorn AP, Goodson JL (2012) Infectious diseases related to travel. In: The Yellow Book. Available: http://wwwnc.cdc.gov/travel/yellowbook/2012/chapter-3-infectious-diseases-related-to-travel/measles-rubeola. Accessed: 14 December 2013.

21. Wallinga J, Heijne JCM, Kretzschmar M (2005) A measles epidemic threshold in a highly vaccinated population. PLoS Med 2(11): e316

22. Hooker G, Ellner SP, Roditi LvD, Earn DJ (2011) Parameterizing state-space models for infectious disease dynamics by generalized profiling: measles in Ontario, JR S Interface 8:961-975.

23. Edmunds WJ, Gay NJ, Kretzschmar M, Pebody RG, Wachmann H. European Seroepidemiology Network. The pre-vaccination epidemiology of measles, mumps and rubella in Europe: implications for modelling studies. Epidemiol Infect 2000; 125: 635-50.

24. Box G E, Draper NR (1987). Empirical model-building and response surfaces. Boston: John Wiley & Sons.

